# Interactive analysis and quality assessment of single-cell copy-number variations

**DOI:** 10.1101/011346

**Authors:** Tyler Garvin, Robert Aboukhalil, Jude Kendall, Timour Baslan, Gurinder S. Atwal, James Hicks, Michael Wigler, Michael C. Schatz

## Abstract

We present an open-source visual-analytics web platform, Ginkgo (http://qb.cshl.edu/ginkgo), for the interactive analysis and quality assessment of single-cell copy-number alterations. Ginkgo automatically constructs copy-number profiles of individual cells from mapped reads, as well as constructing phylogenetic trees of related cells. We validate Ginkgo by reproducing the results of five major studies and examine the data characteristics of three commonly used single-cell amplification techniques to conclude DOP-PCR to be the most consistent for CNV analysis.

---

Single-cell sequencing [1] has become an important tool for probing cancer [2], neurobiology [3], developmental biology [4–6], and other complex systems. Studying genomic variation at the single-cell level allows investigators to unravel the genetic heterogeneity within a sample and enables the phylogenetic reconstruction of subpopulations beyond what is possible with standard bulk sequencing, which averages signals over millions of cells. To date, thousands of individual human cells have been profiled to map the subclonal populations within cancerous tumors [7] and circulating tumor cells [8], to discover mosaic copy-number variations in neurons [3], and to identify recombination events within gametes [5, 9], among many other applications. One key application of single-cell sequencing is to identify large-scale (>10kb) copy-number variations (CNVs) [3, 7, 10]. For example, in cancer, CNVs form a “genetic fingerprint” from which one can infer the phylogenetic history of a tumor [11] and trace progression of metastatic events [7].

Given the insights made possible by single-cell sequencing, many researchers are now interested in applying the technology to study diverse biological systems and species. However, the downstream analysis is complex. Although many approaches and computational tools exist for CNV analysis of bulk samples [12] there are currently no fully automated and open-source tools that address the unique challenges of single-cell sequencing data: (1) extremely low depth of sequencing coverage (< 1X) makes for noisy profiles and makes split-read, paired-end, or SNP density approaches ineffective; (2) amplification biases from WGA markedly distort read counts, including failure to amplify entire segments [13,14]; (3) badly assembled regions of the genome (e.g. centromeres) lead to the artificial inflation of read counts (“bad bins”) [13]; (4) calling copy number at single-cell, integer levels requires development of new algorithms; and (5) exploring population structure is not needed, and often not possible, in bulk sequencing. In addition, several unique sources of cell-specific errors are introduced during the experimental procedures, including GC content and other sequencing biases. While ad hoc methods have been developed for individual studies, there is currently no easy-to-use, open-source software available that executes this pipeline automatically and correctly.

Here we present our new open-source web analytics platform, Ginkgo, for the automated and interactive analysis of single-cell copy-number variations. Ginkgo enables researchers to upload samples, select processing parameters, and after processing, explore the population structure and cell-specific variants revealed within a visual analytics framework in their web browser.

Ginkgo guides users through every aspect of the analysis in a user-friendly interface, from binning reads into regions across the genome, to quality assessment, GC bias correction, segmentation, copy-number calling, visualization and exploration of results (Figure 1). This pipeline builds on our previous single-cell sequencing work [13], and includes several novel features not previously described to advance the state of the art, including: (1) a new algorithm for determining absolute copy-number state from the segmented raw read depth, (2) a new method for controlling quality issues in the reference assembly (see “bad bins” in **online methods**); (3) an option to integrate ploidy information from FACS to accurately call copy number; and (4) a suite of interactive visual analytics tools to allow users to easily share results with collaborators and clinicians. Ginkgo provides functionality for five different species (human, chimp, mouse, rat, and fly) and includes a wide array of tunable parameters for individual users’ needs (**Online Methods**).

**Figure 1:**
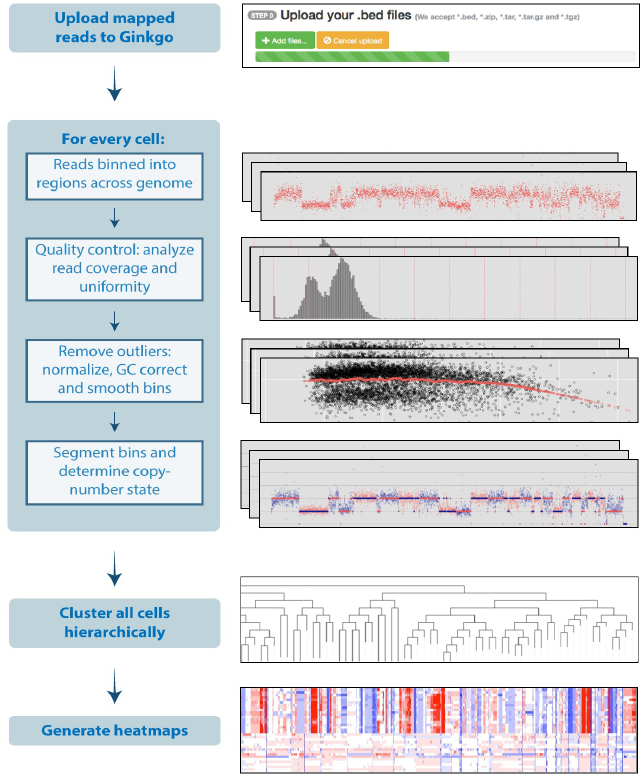
The Ginkgo flowchart for performing single-cell copy-number analysis. Usage and parameters are described in the online methods and on the website.

Once an analysis completes, Ginkgo displays an overview of the data in a sortable data table, an interactive phylogenetic tree [14] of all cells used in the analysis, and a set of heat maps detailing the CNVs that drove the clustering results. Clicking on a cell in the interactive phylogenetic tree or data table allows the user to view an interactive plot of the genome-wide copy-number profile of that cell, search for genes of interest, and link out to a custom track of amplifications/deletions in the UCSC genome browser [26]. Ginkgo also outputs several quality assessment graphs for each cell: a plot of read distribution across the genome, a histogram of read count frequency per bin, and a Lorenz curve to assess coverage uniformity [15]. Subsets of interesting cells can also be selected by the user to directly compare copy-number profiles, Lorenz curves, GC bias, and coverage dispersion.

To validate Ginkgo, we set out to reproduce the major findings of five single-cell studies that used either MALBAC or DOP-PCR amplification. These datasets address vastly different scientific questions, were collected from a variety of tissue types, and make use of different experimental and computational approaches at different institutions. Using Ginkgo, we replicated the published CNVs for each cell in each of the datasets with the exception of one cell in Hou *et al.*, which we believe was due to mislabeling in the NCBI SRA. Moreover, the Navin *et al.* and Ni *et al.* datasets used the identified CNVs to generate phylogenetic trees across all samples. Ginkgo is able to reproduce the distinct clonal subpopulations in the two Navin *et al.* datasets (**Supplementary Figure 1**) and the patient clustering results from Ni *et al.* (**Supplementary Figure 2**). Using simulated copy-number profiles we confirm that Ginkgo reliably identifies copy-number changes (98.8% accuracy, 98.7% true positive rate, 1.2% false positive rate) and perfectly reproduces the population structure through clustering of the individual samples (**Online Methods**).

While Ginkgo corrects for many of the biases present in single-cell data, higher quality data inevitably leads to higher quality results. In order to explore the effects of WGA on data quality, we set out to compare the biases and differences in coverage uniformity between the three most widely published WGA techniques: MDA, MALBAC, and DOP-PCR using 9 distinct datasets, 3 for each method.

Raw sequencing reads from each of nine datasets were downloaded from NCBI (**Online Methods**). Reads were mapped to the human genome and downsampled to match the lowest coverage sample. Finally, aligned reads were binned into 500kb variable-length intervals across the genome such that the intervals average 500kb in length but contain the same number of uniquely mappable positions (see **Online Methods**). We use these binned read counts to measure two key data quality metrics: GC bias and coverage dispersion. Importantly, raw bin counts provide a robust view of the data quality impartial to the different approaches to segmentation, copy-number calling, or clustering.

GC content bias refers to preferential amplification and sequencing because of the percentage of G+C nucleotides in a given region of the genome [16]. This introduces cell-specific and library-specific correlations between GC content and bin counts. In particular, when GC content in a genomic region falls outside a certain range (typically <0.4 or >0.6), read counts rapidly decrease (**Online Methods**). We find that MDA has very high GC bias compared to MALBAC and DOP-PCR (Figure 2a). Only 45.9% of MDA bin counts fall within the expected coverage range compared to 94.0% of MALBAC bin counts and 99.6% of DOP-PCR bin counts. It is important to note that, regardless of WGA approach, each cell has unique GC biases that must be individually corrected.

**Figure 2a.**
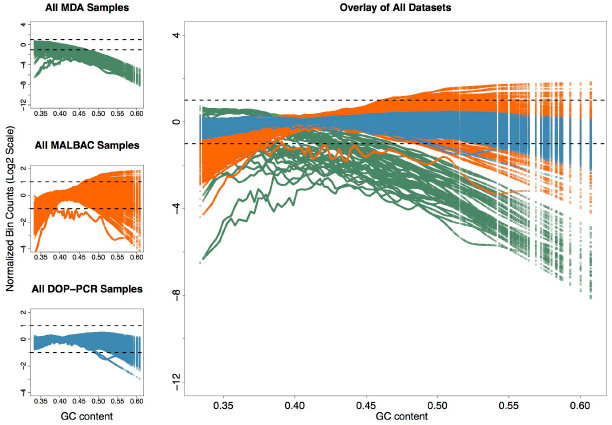
Lowess fit of GC content with respect to log normalized bin counts for all samples in each of the 9 datasets analyzed: 3 for MDA (top left – green), 3 for MALBAC (center left – orange), and 3 for DOP-PCR (bottom left – blue). Each colored line within a plot corresponds to the lowess fit of a single sample. The dashed lines show a two fold increase or decrease in average observed coverage. Note that the three MDA datasets (top left) have a different y-axis scale due to the large GC biases present.

As a further measure of data quality, we calculated the median absolute deviation (MAD) of all pair-wise differences in read counts between neighboring bins for each sample, after normalizing the cells by dividing the count in each bin by the mean read count across bins. MAD is resilient to outliers caused by copy-number breakpoints, as transitions from one copy-number state to another are relatively infrequent. Instead, pair-wise MAD reflects the bin count dispersion due to technical noise. For each of the nine datasets, the MAD was calculated for each cell and displayed in a box-and-whisker plot (Figure 2b). As expected from previous comparisons of MDA to other WGA techniques [15, 17], MDA data displays high levels of coverage dispersion on average, with a mean MAD 2 to 4 times that of the DOP-PCR datasets. In addition, the MALBAC and MDA datasets show large differences in data quality between studies while the DOP-PCR datasets show consistent flat MAD across all three studies (**Supplementary Figure 3**).

**Figure 2b.**
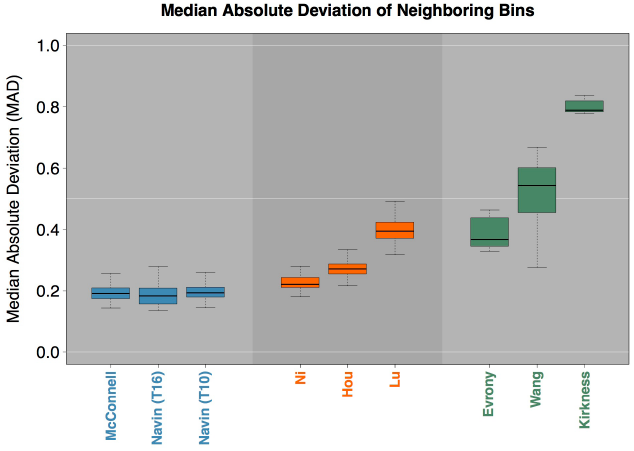
The median absolute deviation (MAD) of neighboring bins: A single pair-wise MAD value is generated for each sample in a given dataset and represented by a box and whisker plot. The high biases present in the MDA datasets make comparing DOP-PCR and MABLAC samples difficult. Figure 1 of the Online Methods shows this comparison more clearly.

We find that DOP-PCR outperforms both MALBAC and MDA in terms of data quality. As previously reported [15, 17–20], MDA displays poor coverage uniformity and low signal-to-noise ratios. Coupled with overwhelming GC biases, MDA is unreliable for accurately determining CNVs compared to the other two techniques. Furthermore, while both DOP-PCR and MALBAC data can be used to generate CNV profiles and identify large variants, DOP-PCR data has substantially lower coverage dispersion and smaller GC biases when compared to MALBAC data. Given the same level of coverage, our results indicate that data prepared using DOP-PCR can reliably call CNVs at higher resolution with better signal-to-noise ratios, and is more reliable for accurate copy-number calls.

## METHODS

Methods and any associated references are available in the online version of the paper.

### Accession codes.

Details are available in Table 2 of the Online Methods.

## Acknowledgements

We would like to thank Nicholas Navin and Peter Andrews for their helpful discussions and assisting getting access to the data. The project was supported in part by National Institutes of Health award (R01-HG006677) to MCS, the National Science Foundation (DBI-1350041) to MCS, the CSHL Cancer Center Support Grant (5P30CA045508), and by the Watson School of Biological Sciences through a Training Grant (5T32GM065094) from the National Institutes of Health.

## Author Contributions

T.G. and R.A. developed the software and conducted the computational experiments. M.C.S, M.W., J.H., and G.S.A. designed the experiments. T.B. and J.K. assisted with the analysis and helped design the experiments. All of the authors wrote and edited the manuscript. All of the authors have read and approved the final manuscript.

