## Supplementary material for "Interactive analysis and quality assessment of single-cell copy-number variations": Online Methods

### 1. Data analysis pipeline and parameters

Ginkgo provides a large number of user specified parameters to control how the analysis is performed and how the results are interpreted. Several of these must be set according to the experimental design of the study (genome, bin size, sex chromosome masking, FACS copy number estimation), while others allow the researcher to explore the analysis using different metrics depending on the goals of the study. See the sections below for a more complete description.

| Category | Available Options |
|---|---|
| Genome | - Human *(hg19, hg18)*<br>- Chimpanzee *(panTro4, panTro4)*<br>- Mouse *(mm10,mm9)*<br>- Rat *(rn5)*<br>- Drosophila *(dm3)* |
| Binning method | - Variable-size bins<br>- Fixed-size bins |
| Bin sizes | - 10kb<br>- 25kb<br>- 50kb<br>- 100kb<br>- 175 kb<br>- 250kb<br>- 500kb |
| Segmentation method | - Independent *(using normalized read counts)*<br>- Global *(using sample with lowest IOD)*<br>- Custom *(user specifies segmentation .bed file)* |
| Clustering method | - Single<br>- Complete<br>- Average<br>- Ward<br>- Neighbor joining |
| Clustering distance metric | - Euclidian<br>- Maximum<br>- Manhattan<br>- Canberra<br>- Binary<br>- Minkowski |
| Masking "bad bins" | - Due to incomplete/incorrect reference assemblies a small number of bins near telomeric and centromeric regions show consistently inflated copy number states across all cells, healthy or otherwise. This option removes these artificial amplifications from the analysis. *Currently only available for human. |

| | |
|---|---|
| Sex chromosome masking | - Binary option (whether or not to mask sex chromosomes during clustering) |
| FACS copy number estimation | - User uploads a file with ploidy estimates from FACS results that are used in tandem with our copy number estimation algorithm (not required). |

**Table 1:** Data processing options in Ginkgo

### 1.a. Binning method

Copy number analysis begins with binning uniquely mapping reads into fixed-length or variable-length intervals across the genome. This aggregates read depth information into larger regions that are more robust to variable amplification and other biases. As discussed in the main text, fixed-length bins are generally discouraged as they lead to read drop out in regions that span highly repetitive regions, centromeres, and other complex genomic regions.

To generate boundaries for variable-length bins, we use the method outlined in [1], where we sample 101bp stretches of the reference assembly at every position along the genome. These simulated reads are mapped back to the genome using Bowtie and only uniquely mapping reads are analyzed. For a given bin size, we assign reads into bins such that each bin has the same number of uniquely-mappable reads. Consequently, intervals with higher repeat content and low mappability will be larger than intervals with highly mappable sequences, although they will both have the same number of uniquely mappable positions.

Using variable-length bins with sufficient depth of coverage and consistent ploidy, sequence reads are expected to map evenly across the entire genome with uniform variance. Users are provided with a variety of bin sizes from which to choose, depending on the overall coverage available; if the mean coverage per bin is too low, we encourage users to use larger bins.

### 1.b. Masking bad bins

As described in the main text, there are a number of regions, specifically around the centromeres of certain chromosomes, where there is an accumulation of very high read depth compared to the expected depth. These regions consistently display high read depth in both bulk and single-cell sequencing data. Using data from 54 normal individual diploid cells from multiple individuals (not presented here), these bins (designated as "bad bins") were determined in the human reference genome (hg19) as follows. The bin counts were divided by the mean bin counts for each cell to normalize for differences between cells in total read count. For each chromosome, the mean of the bins over all cells is subtracted from each individual cell's normalized bin count to normalize for differences between chromosomes. The mean and standard deviation of the autosomes is then used to compute an outlier threshold corresponding to a p-value of 1/N, where N is the number of bins used. In practice, less than 1% of bins are identified as extreme outliers and masked for further processing.

#### 1.c. GC bias correction

Once reads are placed into bins, Ginkgo normalizes each sample and corrects for GC biases prior to segmentation. The normalization process begins by dividing the count in each bin by the mean read count across all bins. This centers the bin counts of all samples at 1.0. To identify and correct GC biases, Ginkgo computes a locally-weighted linear regression using the R function *lowess* [2] (smoother span = .5, iterations = 3, delta=0.1*range(x)) to model the relationship between GC content and log-normalized bin counts. This lowess fit is then used to scale each bin such that the expected average log-normalized bin count across all GC values is zero. After the lowess fit, we monitor the bias of each cell by calculating the proportion of bins that fall outside an expected coverage of zero by +/- 1, log base 2.

#### 1.d. Segmentation (CBS)

Following GC bias correction, bin counts are segmented to reduce fluctuations in noise across chromosomes and identify longer regions of equal copy number. To this end, Ginkgo makes use of Circular Binary Segmentation (CBS) [3], which segments the genome by recursively splitting the chromosomes into segments based on a maximum t-statistic until a reference distribution estimated by permutation is reached. Once the CBS segmentation is complete, the breakpoints (segment boundaries) across all bins are determined, and the counts for all bins within each segment are reset to be the median bin count value within that segment.

The key step during segmentation is selecting the right reference sample for comparison. Using a diploid sample to normalize bin counts can eliminate additional biases uncorrected by GC normalization. Although Ginkgo supports uploading data from such a cell, this is not always available so Ginkgo provides alternatives for segmenting samples: (1) Independent segmentation, where samples are segmented independently by their own normalized bin count profiles; and (2) Sample with lowest IOD, where Ginkgo selects the sample with the lowest index of dispersion (IOD - the ratio between the read coverage variance and the mean) and uses that sample as a reference for all other samples. The sample with the lowest index of dispersion will likely be among the most evenly balanced ploidy and highest quality of all submitted cells.

#### 1.e. Determining copy-number state

Since we are analyzing single-cell data, we expect every genomic locus to have an integer copy number (CN) value. Furthermore, the quantized nature of single-cell data means that the same number of reads per bin should separate every sequential CN state, e.g., ~50 reads for CN 1, ~100 reads for CN 2, ~150 reads for CN 3, etc. While biological and technical noise prevent read counts from segregating perfectly into distinct CN states, read counts should still be centered around integer CN states.

The most direct approach for determining the CN state of each cell is available for users that have *a priori* knowledge of the ploidy of each sample. For example, cells that are DAPI-stained prior to cell sorting can be gated based on their fluorescence activity, and ploidy can be determined by comparing its fluorescence activity to that of a reference cell with a known CN state. With these data, Ginkgo determines the copy number state of each sample by scaling the segmented bin counts such that the mean bin count is equal to the ploidy of the sample. Finally bin counts are rounded to integer copy number values. Advances in fluorescence activated cell sorting (FACS) will make this copy number prediction even more accurate in time, although cells that are incorrectly sorted and placed into wells with more than one cell will show much higher fluorescence activity and will have an incorrectly inferred copy number state.

Since FACS data is not always available for analysis and has potential for error, Ginkgo provides an alternative to determine the copy number of each sample. As discussed earlier, before determining the CN state of a cell, the cell is binned, normalized, and segmented. This copy number profile has a mean of one and is referred to as the raw copy number profile (RCNP). If the true genome-wide copy number of a sample were equal to X, the scaled copy number profile (SCNP) would then be the product of RCNP and X, and the final *integer* copy number profile (FCNP) would be the rounded value of the SCNP so all segments contain an integer value.

With these relationships, Ginkgo infers the genome-wide copy number X using numerical optimization. For a given cell, Ginkgo first determines the SCNP and FCNP for all possible values of X in the set [1.50, 1.55, 1.60, … , 5.90, 5.95, 6.00]. Ginkgo then computes the sum of square (SoS) error between the SCNP and the RCNP for each value of X and selects the value of X with the smallest SoS error. Once the multiplier is identified and applied, the scaled bins are rounded to generate the final integer copy number profile for each sample. Intuitively, this is equivalent to finding the copy number multiplier that causes the normalized segmented bin counts to best align with integer copy number values.

#### 1.f. Clustering

Before visualization, the final step is to look outside the scope of individual cells and determine the overall population structure. Ginkgo first determines the distance (dissimilarity structure) between all cells. We provide six choices of distance metrics: Euclidean, maximum, Manhattan, Canberra, binary, and Minkowski. After computing the dissimilarity matrix, Ginkgo then computes a dendrogram through neighbor joining or by hierarchically clustering samples using one of four different agglomeration methods: single linkage, complete linkage, average linkage, and ward linkage. In addition, Ginkgo generates a phylogenetic tree by first computing the Pearson correlation between all samples and using these dissimilarity values to cluster the samples.

#### 1.g. Masking sex chromosomes

Careful consideration of gender must be given when analyzing patients from mixed populations, as the combined set of the X and Y-chromosomes make up a large fraction of the human genome that can distort the clustering results. Indeed, when we examined the Ni *et al*. dataset with Ginkgo with sex chromosomes masked, we could still discriminate between individual patient's tumors, but we could no longer discriminate between ADC and SCLC (**Supplementary Figure 2B**); the SCLC patients were exclusively female and, with one exception, the ADC patients were entirely male. Ginkgo comes prepackaged with the ability to selectively mask sex chromosomes to prevent gender biases from dominating the clustering.

### 2. Single-cell datasets analyzed

We validate Ginkgo by reproducing major findings of several single cell sequencing studies that employ three different WGA techniques: MALBAC, DOP-PCR/WGA4, and MDA. Take together, we analyze the data characteristics of nine datasets across five tissue types (Table 2). The Ginkgo parameters for these datasets are described in the main text, and additional parameters are noted below.

| Study | WGA Method | Tissue Type | # of cells | Accession |
|---|---|---|---|---|
| Kirkness *et al.* (2013) | MDA | Sperm | 11 | SRP017516 |
| Wang *et al.* (2012) | MDA | Sperm | 31 | SRA053375 |
| Evrony *et al.* (2012) | MDA | Neuron | 8 | SRA056303 |
| Lu *et al.* (2012) | MALBAC | Sperm | 99 | SRA060945 |
| Ni *et al.* (2013) | MALBAC | Lung | 29 | SRP029757 |
| Hou *et al.* (2013) | MALBAC | Oocyte | 181 | SRA091188 |
| Navin *et al.* (2011) | DOP-PCR | Breast (T10) | 100 | SRX021401 |
| Navin *et al.* (2011) | DOP-PCR | Breast (T16) | 100 | SRX037035/SRX037132 |
| McConnell *et al.* (2013) | DOP-PCR | Neuron | 109 | SRP030642 |

**Table 2:** List of the 9 datasets analyzed across 8 different studies.
Note that there are two distinct datasets from the same Navin *et al.* study.

Reads were mapped to hg19 using bowtie [4] and only uniquely mapped reads (mapping quality score >= 25) were kept. Mapped read counts ranged from 1,538,234 (Ni et al.) to 30,638,853 (Lu et al.) with a mean of 15,827,886. To perform an unbiased comparison, all samples were randomly downsampled to 1,538,234 reads to match the lowest available coverage.

In order to compute the GC biases across all nine datasets we calculate the lowess fit of the log base 2 normalized read counts with respect to the bin GC content for each sample. A sample with no GC bias would have a flat normalized read count of zero across all bins and all GC values. After the lowess fit, we monitor the bias of each cell by calculating the proportion of bins that show a two fold change from the expected coverage in either direction (by +/- 1, log base 2).

### 3. Detailed comparison of MALBAC and DOP-PCR protocols

Whole-genome amplification using MDA introduces a large degree of biases compared to MALBAC or DOP-PCR, limiting its applicability to CNV analysis. As such, we focused the scope of the remaining comparisons on the latter two WGA techniques. For a fine-grained comparison of MALBAC and DOP-PCR, we compare the T10 dataset from Navin *et al*. and the CTC dataset from Ni *et al*. due to their similar biological and technical conditions and similar published analysis. Both datasets contain aneuploid cancer cells, were sequenced to similar depth (CTC mean read count: 4,133,466; T10 mean read count: 6,706,119), and were used to generate phylogenetic clusters of samples based on CNVs. We begin by comparing the coverage dispersion and investigate the minimum coverage and bin size needed to reproduce the published results.

#### 3.1. Coverage dispersion

Using the MAD criteria described above, the DOP-PCR-based T10 dataset shows markedly better bin-to-bin correlation than the MALBAC-based CTC dataset as judged by a lower MAD of adjacent and offset bin counts (**Figure 1**). For adjacent bins, the first quartile of the CTC MAD comparison (orange) is higher than the third quartile of the T10 MAD comparison (blue). As we increase the bin offset, greater variation is seen in the CTC data as show by the separation of the mean MAD between the T10 and CTC datasets. We interpret this to mean that there is more local trending in amplification efficiency in MALBAC than in DOP-PCR data.

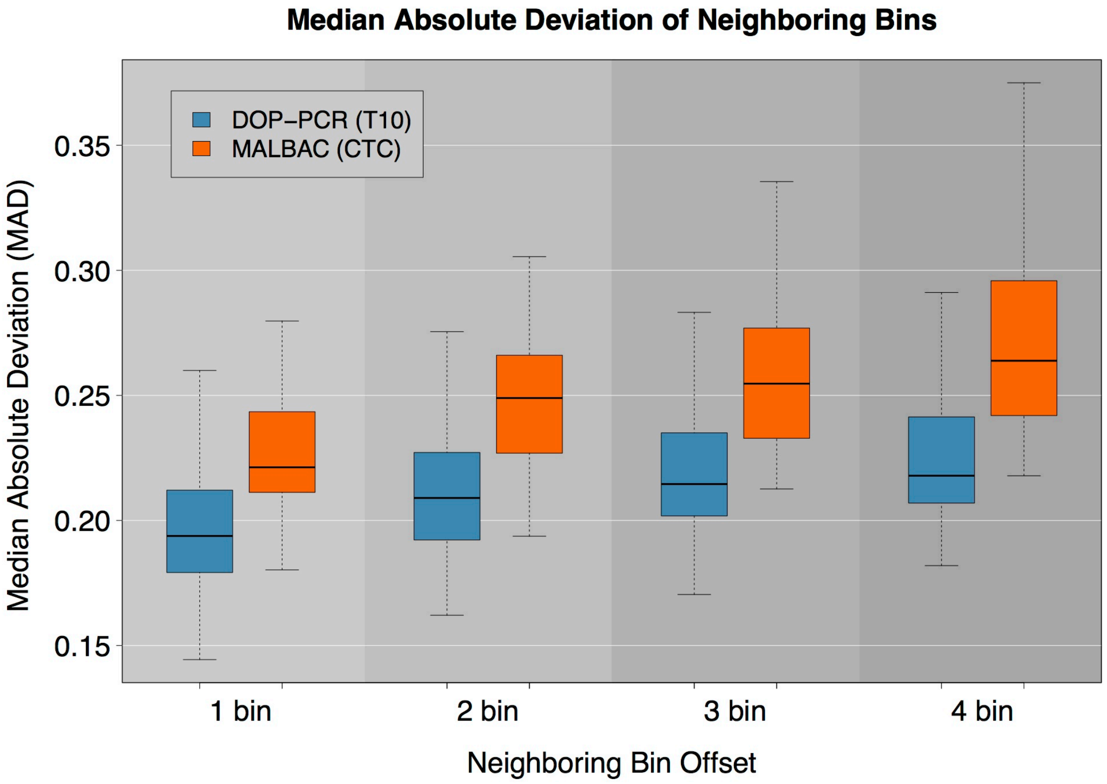


**Figure 1 |** A comparison of MAD between the Navin *et al*. (T10) shown in blue and Ni *et al*. (CTC) shown in orange. As the bin offset increases the separation between the mean T10 MAD and mean CTC MAD grows.

### 3.2. Minimum coverage requirement

We next explore whether WGA protocols differ with respect to the minimum coverage required to observe the same population/clonal substructure identified at full coverage. To this end, we down-sample all datasets and analyze each in Ginkgo to determine: (1) how well segment breakpoints are conserved and (2) how well the phylogenetic relationships are maintained. With all degrees of downsampling (from 25% to 99%), the T10 data shows better breakpoint conservation than the CTC data, but as expected, increased degrees of downsampling lead to substantial erosion of breakpoint boundaries in both datasets (see **Supplementary Figure 4**).

Nevertheless, these downsampling experiments demonstrate MALBAC and DOP-PCR are remarkably robust with respect to preserving the overall clonal/population structure, even at extremely low coverage, although additional smaller CNVs can be discovered with deeper coverage [5]. The clonal structure of the T10 dataset remains fully intact across all downsampling experiments even as the mapped reads are downsampled by 99% (from ~608 reads/bin to ~6 reads/bin). The population structure of the CTC dataset is preserved when downsampled by 95% (from ~597 reads/bin to ~30 reads/bin); when downsampled to 99%, one cell from one patient is incorrectly clustered.

Although depth of coverage in both studies was originally very low (< 0.15x), our downsampling results indicate that Ginkgo can correctly determine the phylogenetic relationship between samples even when sequenced to a depth of coverage of only 0.01x. If generally applicable, which we have not proven here, this approach will allow sparser sequencing with higher throughput at equivalent cost. After low-coverage sequencing, a number of cells from the same phylogenic branch can be pooled for deeper sequencing if desired.

### 3.3. Optimizing bin sizes

Bin size directly impacts the resolution at which CNVs can be called [6]. Thus far we used 500kb-bins to reproduce the results of Navin *et al*. and Ni *et al*. following the procedure by Ni *et al*. However, such large bin sizes hinder the identification of smaller copy-number events. To identify the minimum bin size needed to reproduce the published results, we decreased bin size from 500kb to 10kb (**Table 1)** for both datasets until the hierarchical clustering of the copy number profiles produced different results.

The T10 dataset retained its hierarchical structure until bin sizes dropped below 25kb (**Supplementary Figure 5**), while the CTC dataset lost it original hierarchical structure at a bin size of 100kb. In the T10 dataset, when bin sizes drop to 10 kb, a few hypodiploid cells incorrectly cluster. In the CTC dataset, as bin sizes approach 100kb, cells from two patients (4 and 7) begin to overlap. Using 50kb bins, there is widespread overlap between nearly all patients' cells, and only the cells from two patient cluster correctly (**Supplementary Figure 6**). This indicates that at the same level of coverage, DOP-PCR can resolve smaller CNVs than MALBAC, but more comparably structured studies are needed.

### 3.4 Detecting integer copy-number states

Preliminary analysis of bin counts indicate that at the same level of coverage, MALBAC data had a higher level of coverage dispersion and therefore a worse signal to noise ratio than DOP-PCR data. Our downsampling experiments support this claim as the ability to properly discriminate between CTC patients based on the CN states is lost at a bin resolution that is easily resolved with the T10 dataset. To understand the effects of noise further, we evaluated each dataset to discriminate distinct copy number states.

Because the copy-number states of individual cells are integer, we expect the data to be centered at integer values. If the data is highly uniform, read coverage per bin will tightly surround integer copy-number states. As bin count dispersion around copy-number states increases, or is influenced by local chromosomal trends, the distinction between copy-number states will blur.

To examine this, we generated a histogram of the normalized read count distribution for the CTC and T10 datasets (**Supplementary Figure 7**). **Figure 2** shows distributions of bin counts for representative cells: excellent, typical, and lower quality cells as well as the highest quality population average. All T10 profiles have distinct peaks representative of integer copy-number values. While there are a few cells in the CTC dataset that have distinct peaks, many of the CTC profiles have considerably worse resolution with substantial blurring between CN states. Furthermore, the scaled read count distributions illustrate the substantial difference in signal-to-noise between the T10 and CTC datasets (**Supplementary Figure 8**).

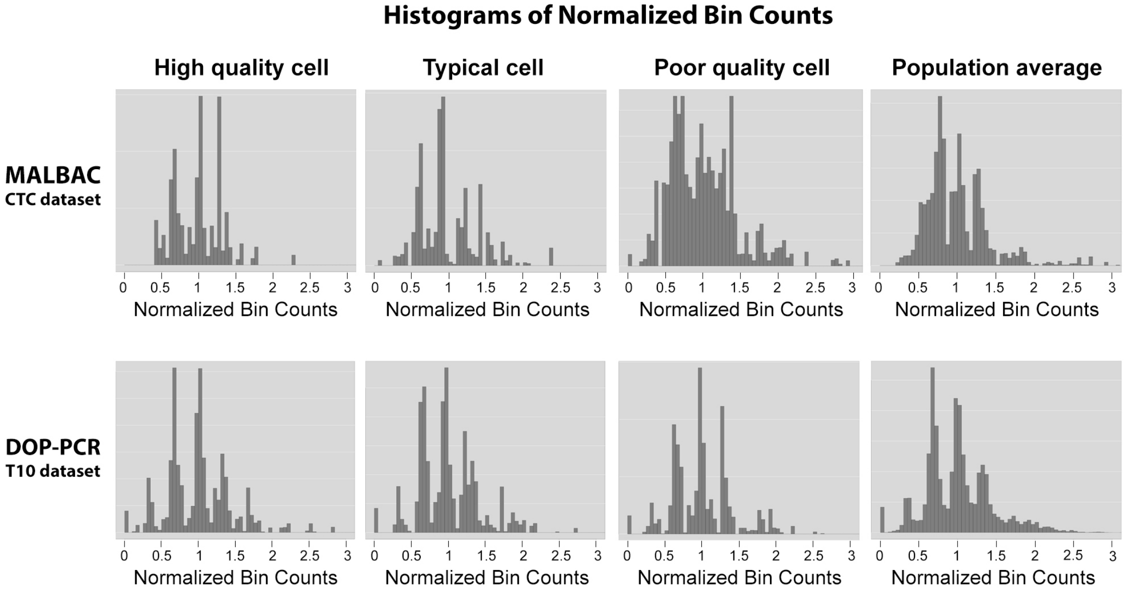


**Figure 2 |** Histograms of normalized bin counts across the CTC and T10 datasets, for a high-, typical-, and poor-quality cell. The rightmost column contains histograms of high quality cell population averages. Distinct peaks are representative of clean data from which accurate copy number calls can be made.

### 4. Simulation analysis of copy number accuracy

To test the accuracy of the copy number and clustering analysis by Ginkgo, we simulated single cell sequencing of 90 cells with 100 total copy-number events per cell. We modeled the cells after a population comprised of 9 distinct clonal populations, with 10 cells per population (**Figure 3a**). We began by generating 3 primary clonal populations by introducing 80 copy-number events compared to the parent diploid cell. Next, for each of the 3 primary clones, we generated 3 subclonal populations by introducing an additional 20 non-overlapping copy-number events to the original clones. Overall, this resulted in 9 distinct subclones belonging to 3 larger clonal populations with a total of 100 CNVs with respect to the human reference genome (hg19).

The genome positions of CNVs were non-overlapping and generated from a uniform random distribution across the genome. The lengths of CNVs were generated from an exponential distribution with a mean of 5Mb and bounded between the range of 200kb and 20Mb to approximate the CNVs observed in the genuine data. The copy-number state of the CNVs were generated from a Poisson distribution with a mean of 2.5 excluding the value 2.

We generated 10 cells from each of the 9 subclones (90 cells in total) by simulating reads from the subclone reference sequences generated above. For each cell, we simulated 200k, 101bp, single-end reads from the subclone reference sequence using dwgsim (https://github.com/nh13/DWGSIM) (dwgsim –n 101 –z -1 –e .01 –d 1 –r 0 -1 101 -2 0). For each cell, the simulated reads were then mapped to the hg19 human reference genome using the command "bowtie hg19.fa –S –t –m --best –strata" and filtered for only uniquely mappable high scoring reads (quality > 25). The SAM output was then converted to BED format and all 90 cells were uploaded and analyzed directly within Ginkgo with variable length 50kb bins.

Ginkgo is able to accurately reproduce the population structure through hierarchical clustering (**Figure 3b**). In addition, we examined Ginkgo's ability to call CNVs by examining the false negative and false positive rates for all 90 cells at three different read counts (2M, 1.5M, 1M) across three different bin sizes (100kb, 50kb, 25kb) (**Supplementary Table 1**). We measured Ginkgo to have a 0.15% negative and 0.08% false positive rate excluding those bins that are partially spanned by a copy number alteration. When the entire genome is considered, including partially spanned bins, Ginkgo still has only an ~2% false negative and ~1.2% positive rate. Hence, as expected, errors are almost exclusively concentrated at the boundaries of CNVs where the precise end of the event cannot be determined due to the extremely low coverage available or partially spanning of a bin.

We compared these results to the widely used CNVnator algorithm [7] for bulk sequencing CNV analysis and find that Ginkgo performs CNV calls with higher accuracy (**Supplementary Table 1**). Furthermore, CNVnator and other bulk sample analysis programs do not attempt to assign integer copy number states, but in this analysis we have measured Ginkgo's accuracy with this more strict requirement while for CNVnator we could only evaluate if an amplification or deletion had been identified. Ginkgo also has numerous features for evaluating population-wide CNV relationships (heatmaps & hierarchical clusters, multi-sample GC & Lorenz plots, etc) that are also not present in CNVnator or other bulk sample programs that we could not evaluate. Finally, in a practical sense, we also find Ginkgo to be substantially faster than CNVnator, requiring a few hours via a simple web-interface rather than many days in a very difficult to install console program for the 90 cell evaluation.

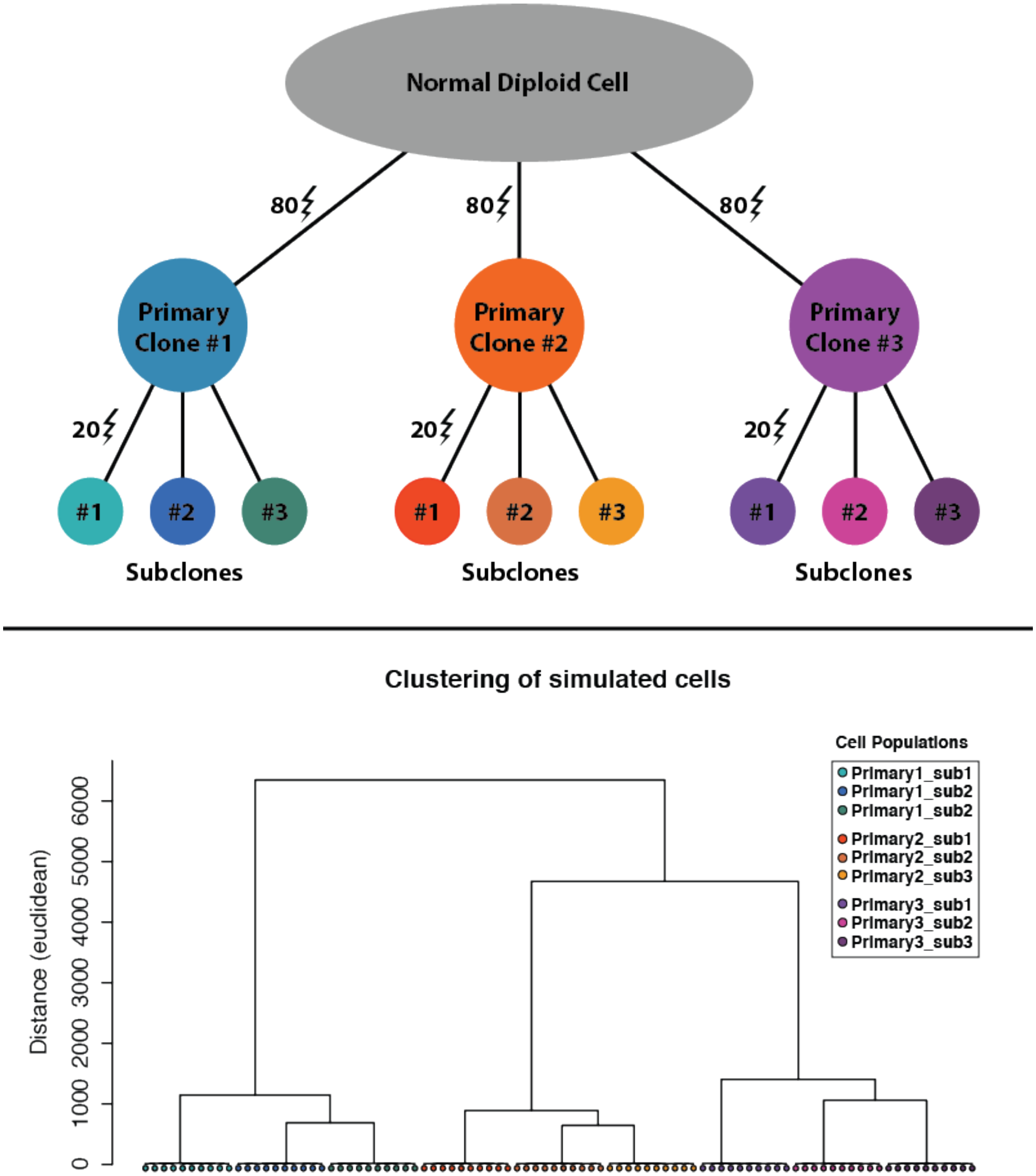


**Figure 3 | (A) Model representation of the 9 distinct subclones generated by simulation of 100 copy number events with respect to the reference. (B) Hierarchical clustering of the 90 samples by Ginkgo. Ginkgo perfectly recovers the underlying subclonal population structure.**
