## Supplementary material for "Interactive analysis and quality assessment of single-cell copy-number variations": Online Methods

**Supplementary Materials**

### Supplementary Note 1: Replication of the single-cell analysis results

#### 1. Navin et al.

This work profiled breast cancer in two separate studies. The first (dataset T10) examined heterogeneity in a polygenomic breast tumor. CNV analysis and hierarchical clustering of 100 single-cells revealed three distinct clonal subpopulations present in the tumor. The second study (datasets T16M/P) examined a monogenomic breast tumor and its suspected liver metastasis. CNV analysis and hierarchical clustering of 100 cells revealed that a single clonal expansion formed the primary breast tumor and seeded the metastasis. In the polygenomic breast tumor analysis, Ginkgo clusters all 100 samples into the same four distinct subpopulations of the original study, replicating the published population structure (**Supplementary Figure 1A**). In the monogenic breast tumor and its associated liver metastasis analysis, Ginkgo clusters all 100 samples into the same three distinct subpopulations as the original publication, linking the primary tumor to its metastasis (**Supplementary Figure 1B**).

#### 2. McConnell et al.

This study profiled CNV events in human hiPSC-derived fibroblasts and 110 frontal cortex neurons. McConnell et al. found a wide degree of mosaic copy-number variation in neurons and discovered that a subset of neurons have highly aberrant genomes. Using Ginkgo, we successfully reproduce the major aneuploid events in all frontal cortex neurons.

### 3. Ni et al.

This study explored SNPs and CNVs in circulating tumor cells (CTCs) in patients with lung cancer. Through CNV analysis and hierarchical clustering of 29 CTCs across 7 patients with lung adenocarcinoma (ADC) or small-cell lung cancer (SCLC) the authors discovered that CNVs appear specific to cancer types and are reproducible from cell to cell and from patient to patient. Using default settings in Ginkgo, we generate similar CN profiles for all 29 samples and can reproduce the published clustering results (**Supplementary Figure 2**).

#### 4. Hou et al.

This study sequenced the triads of first and second polar bodies (PB1 and PB2) and the oocyte pronuclei from same female egg donors to phase these genomes and determine their crossover maps and frequency. Additionally, genome-wide CN profiles were generated to explore aneuploidy in each sample. The authors identified a total of 47 CNVs in 25 aneuploid cells across 5 patients. We could replicate these results as Ginkgo uncovered 45 of the 47 CNVs in 23 of the 25 identified aneuploid cells. One sample, S0808 (containing the missing two cells/CNVs), did not have CN events matching the published results. We believe this was due to accidental mislabeling of samples/sample IDs upon being deposited to NCBI.

**5. Lu et al.**

In this study, single-cell sequencing of 99 sperm cells from an Asian male were used to examine meiotic recombination and aneuploidy. Across the 99 samples, the authors uncovered 5 aneuploid cells. As expected, our CNV analysis and hierarchical clustering using Ginkgo was able to separate the X and Y bearing chromosomes with the exception of two cells with extremely poor coverage uniformity and a high degree of read drop out that clustered separately (**Supplementary Figure 8**). In addition, we could cleanly replicate the CNV results as Ginkgo uncovered the same chromosomal aberrations in the five aneuploid cells as the original study.

**6. Kirkness et al.**

This work profiled genomic variants in sperm with the goal of demonstrating a technique to retrieve complete haplotypes of sequenced genomes, information that is averaged out and lost during conventional genome sequencing approaches. Using genotype data from 16 cells and low coverage single-cell sequencing of 96 sperm cells, the authors leverage the haploid nature of sperm to identify recombination events at a median resolution less than 100 kb.

**7. Wang et al.**

Much like *Kirkness et al.*, this work examined meiotic recombination and de novo mutations in sperm. Using a microfluidic system, the authors carried out single-cell sequencing on 91 sperm cells to generate a personal recombination map. This map was confirmed by the low-resolution afforded by bulk sequencing but individual samples were found to have significant differences from pedigree data at higher resolution. Finally, the authors used these results to test for meiotic drive, gene conversion, and genome instability. Deep sequencing of 8 single-cells revealed additional unique de novo mutations.

**8. Evrony et al.**

This work looked to unravel to what extent genetic mosaicism exists in the brain of an individual. To this end the authors sequenced 300 single neurons from the cerebral cortex and caudate nucleus of three normal individuals. L1 insertion profiling led to estimate that there are less than < 0.6 unique somatic L1 insertions per neuron with most neurons (~80%) lacking any detectable unique somatic mutations. Finally, the authors genotyped single cortical cells of a child with hemimegalencephaly to characterize the mosaicism of a somatic AKT3 mutation. The AKT3 mutation was found in both neuronal and nonneuronal cells indicating that the mutation occurred in nueroglial progenitor

| Simulated reads (millions) | Mapped reads (millions) | Mean bin length (kbp) | False Negative Rate (%) | | | False Positive Rate (%) | | |
|---|---|---|---|---|---|---|---|---|
| | | | Ginkgo Complete | Ginkgo | CNVnator | Ginkgo Complete | Ginkgo | CNVnator |
| 2.0 | 1.64 | 100 | 0.15 | 2.03 | 6.37 | 0.08 | 1.28 | 0.69 |
| 2.0 | 1.64 | 50 | 0.18 | 1.29 | 5.86 | 0.07 | 1.20 | 0.5 |
| 2.0 | 1.64 | 25 | 0.26 | 1.63 | 6.01 | 0.05 | 1.16 | 0.54 |
| 1.5 | 1.23 | 100 | 0.22 | 2.22 | 6.46 | 0.10 | 1.34 | 0.75 |
| 1.5 | 1.23 | 50 | 0.28 | 1.67 | 5.99 | 0.07 | 1.21 | 0.66 |
| 1.5 | 1.23 | 25 | 0.39 | 2.37 | 6.1 | 0.08 | 1.21 | 0.6 |
| 1.0 | 0.82 | 100 | 0.33 | 2.47 | 6.42 | 0.17 | 1.41 | 0.94 |
| 1.0 | 0.82 | 50 | 0.50 | 2.17 | 6.23 | 0.13 | 1.24 | 1.03 |
| 1.0 | 0.82 | 25 | 0.75 | 3.82 | 6.03 | 0.14 | 1.24 | 0.68 |

### Supplementary Table 1: Simulation accuracy.

False negative and false positive rates for genomes with 100 simulated copy number events at varying read depths and bin sizes. “Ginkgo complete” represents only the segments of copy number variants that fully overlap bin boundaries.

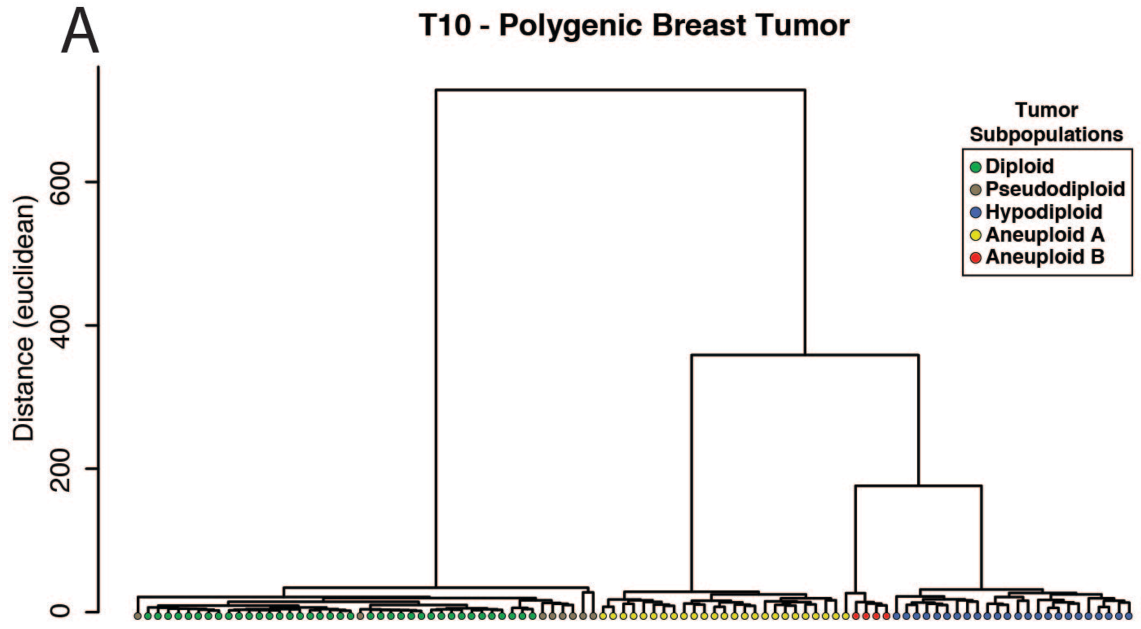

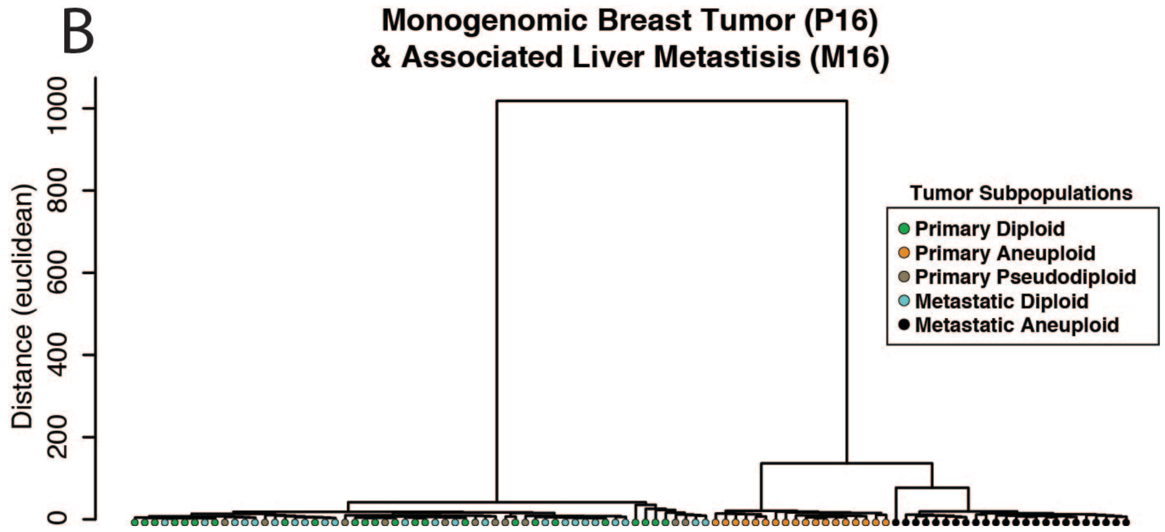

### Supplementary Figure 1: Ginkgo analysis of the Navin et al. cancer data

Phylogenetic trees generated through hierarchical clustering by copy-number using (A) 100 polygenomic breast tumor samples (T10) and (B) 52 monogenomic breast tumor (T16P) and 48 liver metastasis (T16M) samples. These results match the clonal structure published in the original study.

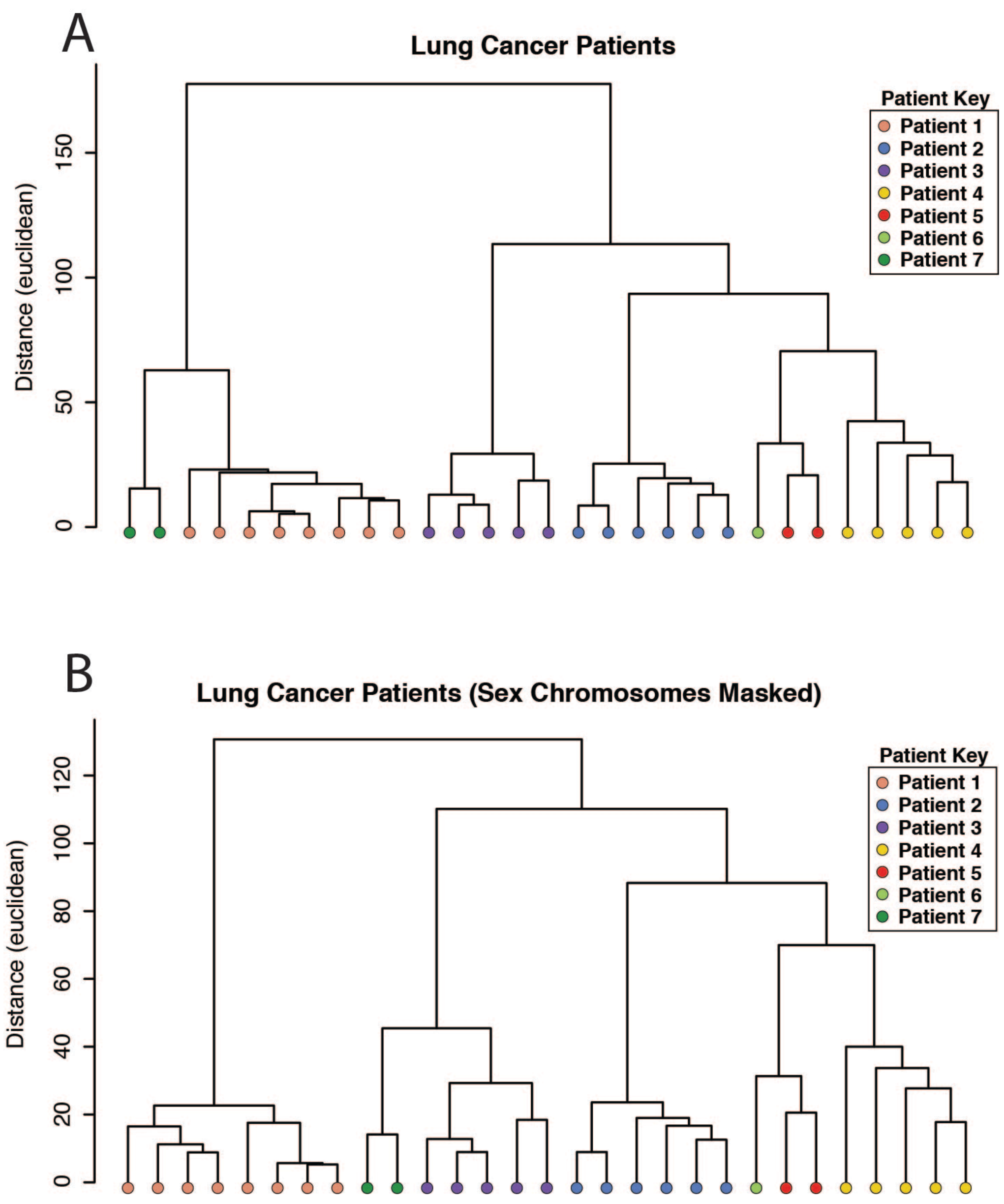

### Supplementary Figure 2: Ginkgo patient clustering of Ni et al. CTC data

(A). Hierarchical clustering by Ginkgo of 29 samples derived from 7 different patients with either adenocarcinoma (patients 2-6) or small cell lung cancer (patients 1, 7), matching the results published by Ni et al. (B) When sex chromosomes are masked, cells still cluster by patient, but patients no longer cluster by cancer subtype. In particular, after masking sex chromosomes, patient 3 is intermixed between patients 1 and 7 and there is no clear association between cancer types.

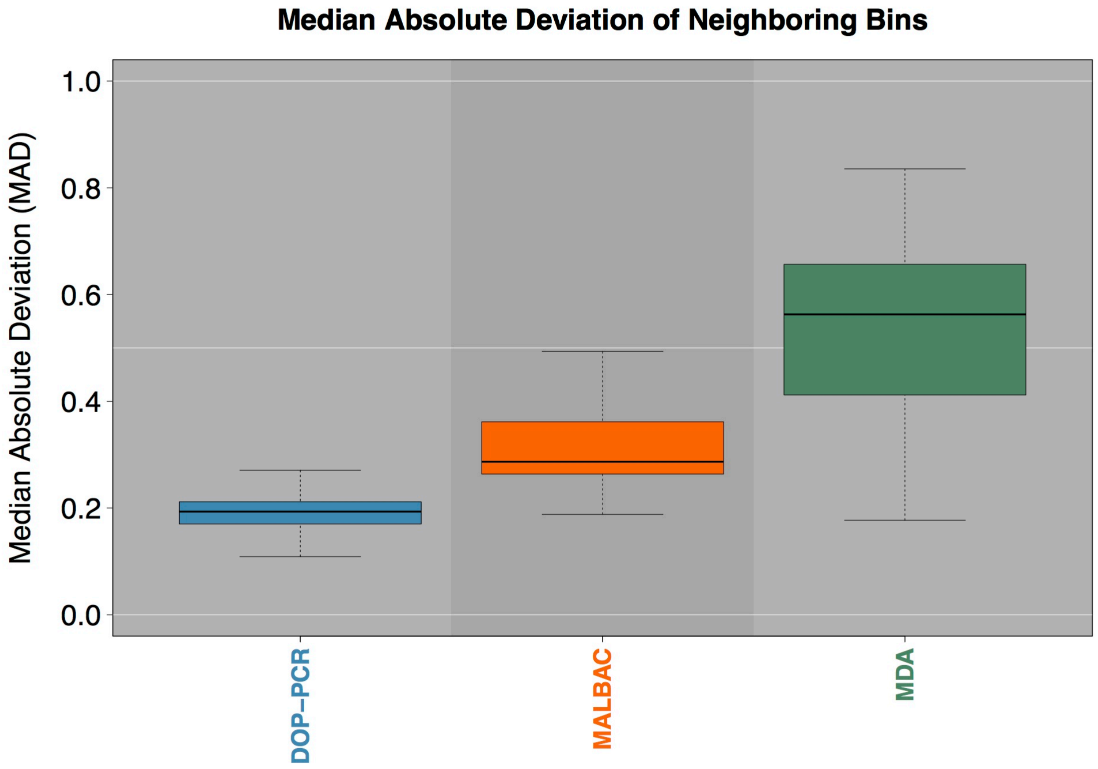

**Supplementary Figure 3: The median absolute deviation (MAD) of neighboring bins with the 3 WGA approaches.**

A single pair-wise MAD value is generated for each sample in a given dataset and represented by a box and whisker plot. The DOP-PCR datasets show the lowest mean MDA as well as the lowest variance across samples. While certain MDA samples outperform the MALBAC dataset, they show much large variability in data quality than MALBAC.

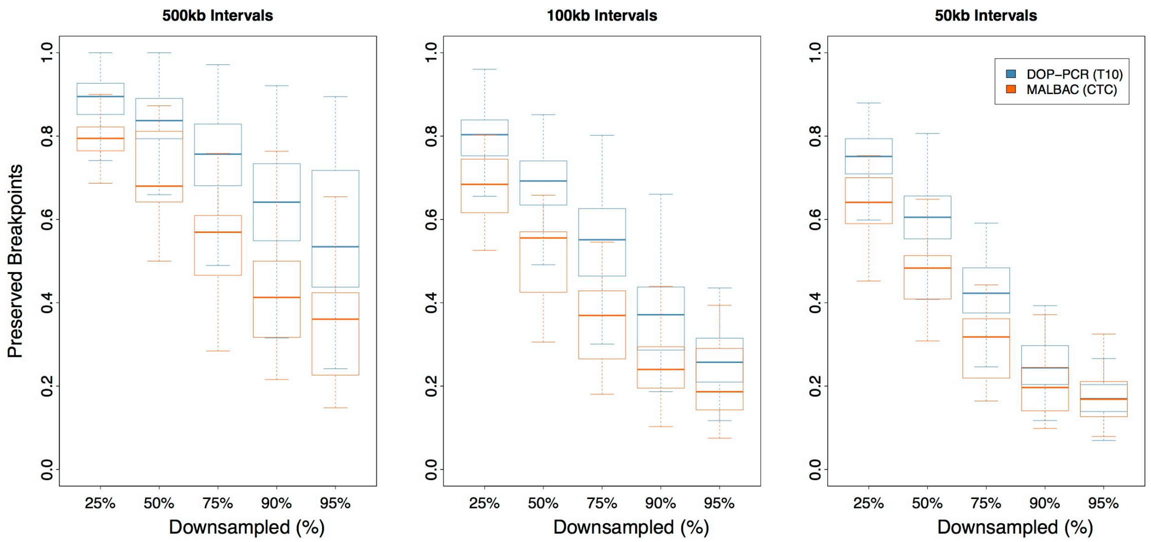

### Supplementary Figure 4: Comparing breakpoint conservation between T10 and CTC.

The fraction of breakpoints conserved between the full intact dataset and the dataset downsampled 25%, 50%, 75%, 90%, or 95% =using interval sizes of **(A)** 500kb, **(B)** 100kb, and **(C)** 50kb. T10 breakpoints are shown in blue and CTC breakpoints are shown in orange. At all levels of downsampling and all intervals sizes, the T10 DOP-PCR data retains a larger fraction of breakpoints than the CTC MALBAC data.

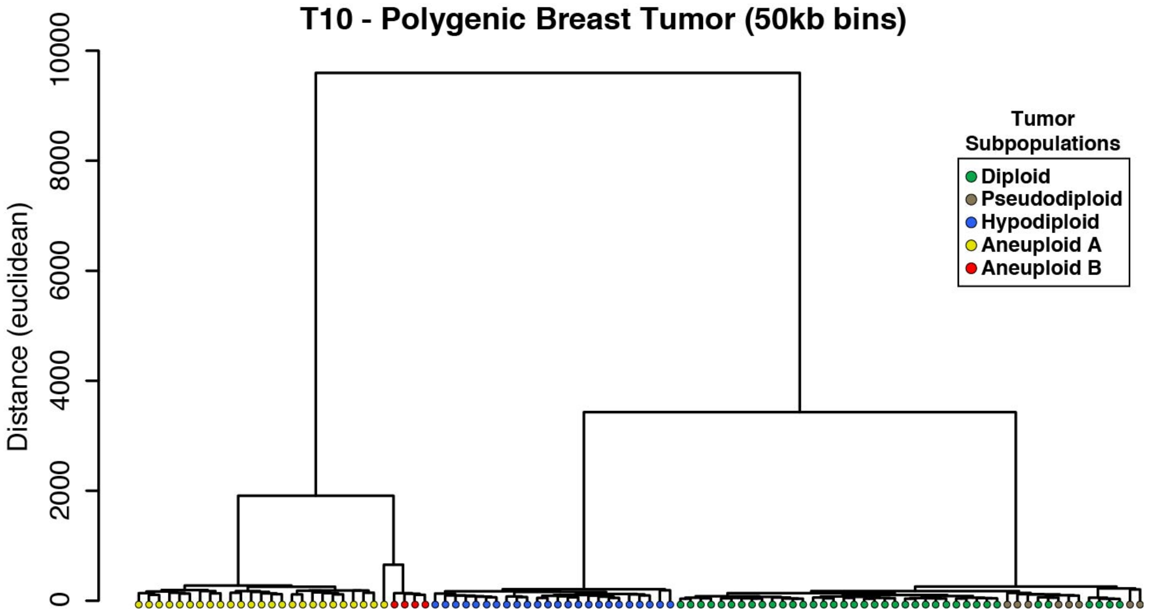

### Supplementary Figure 5: Phylogenetic tree of the T10 cell population

In this analysis, Ginkgo used a bin size of 50kb using the copy number profiles of individual cells. All of the clonal populations remain intact and represent the 500kb clustering results.

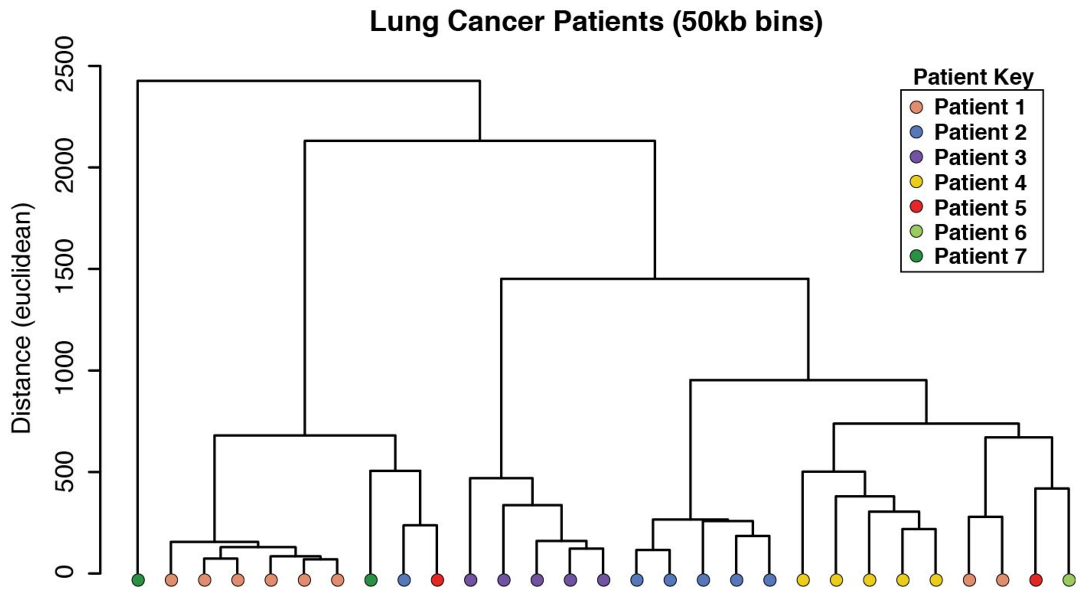

### Supplementary Figure 6: A phylogenetic tree of the CTC single cell samples

In this analysis, Ginkgo analyzed all 7 patients at a bin size of 50kb using the copy number profiles of the individual cells. Only cells from patients 2 and 4 can be correctly clustered at this bin resolution. The cells of the remaining patients no cluster together.

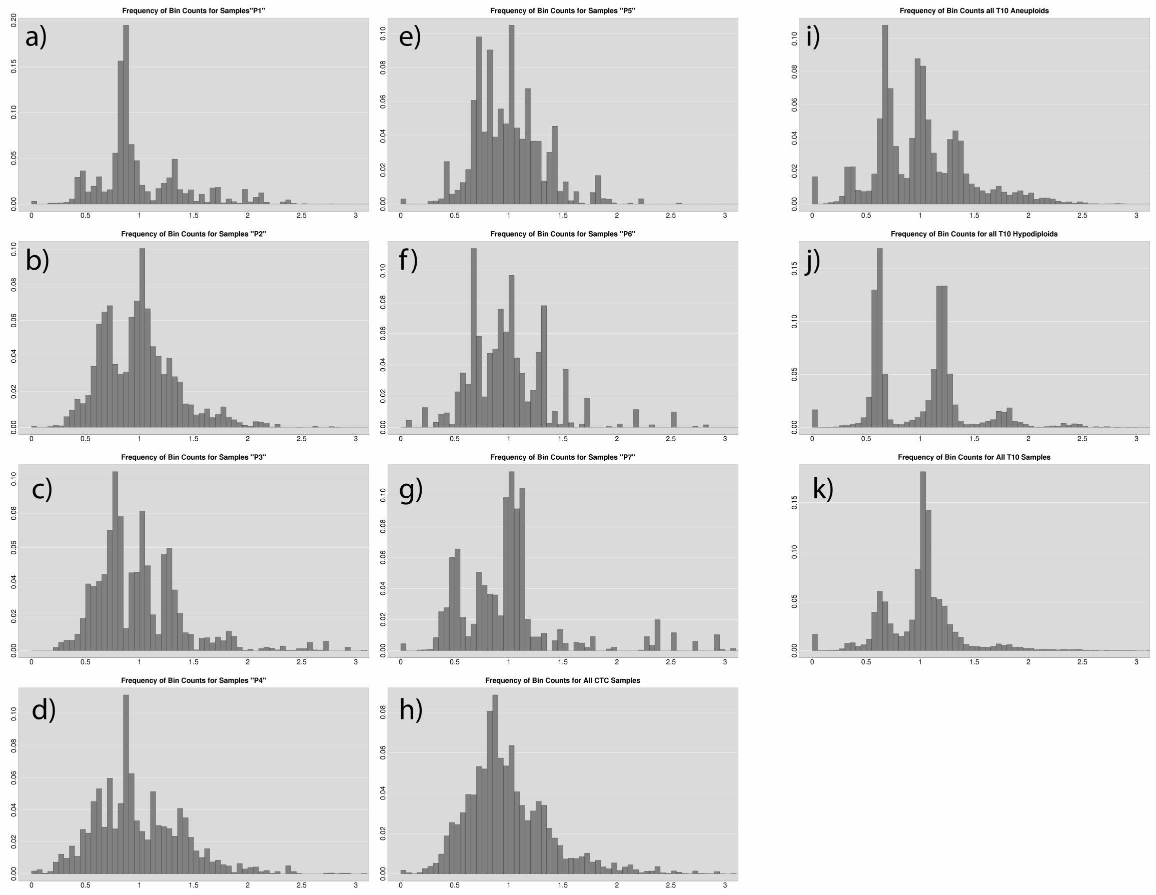

### Supplementary Figure 7: Histograms of normalized bin counts

This analysis was performed across **(a-g)** patients 1-7 respectively, **(h)** all CTC patients, **(i)** all T10 hypodiploid samples, **(j)** all T10 aneuploid samples, and **(k)** all T10 samples.

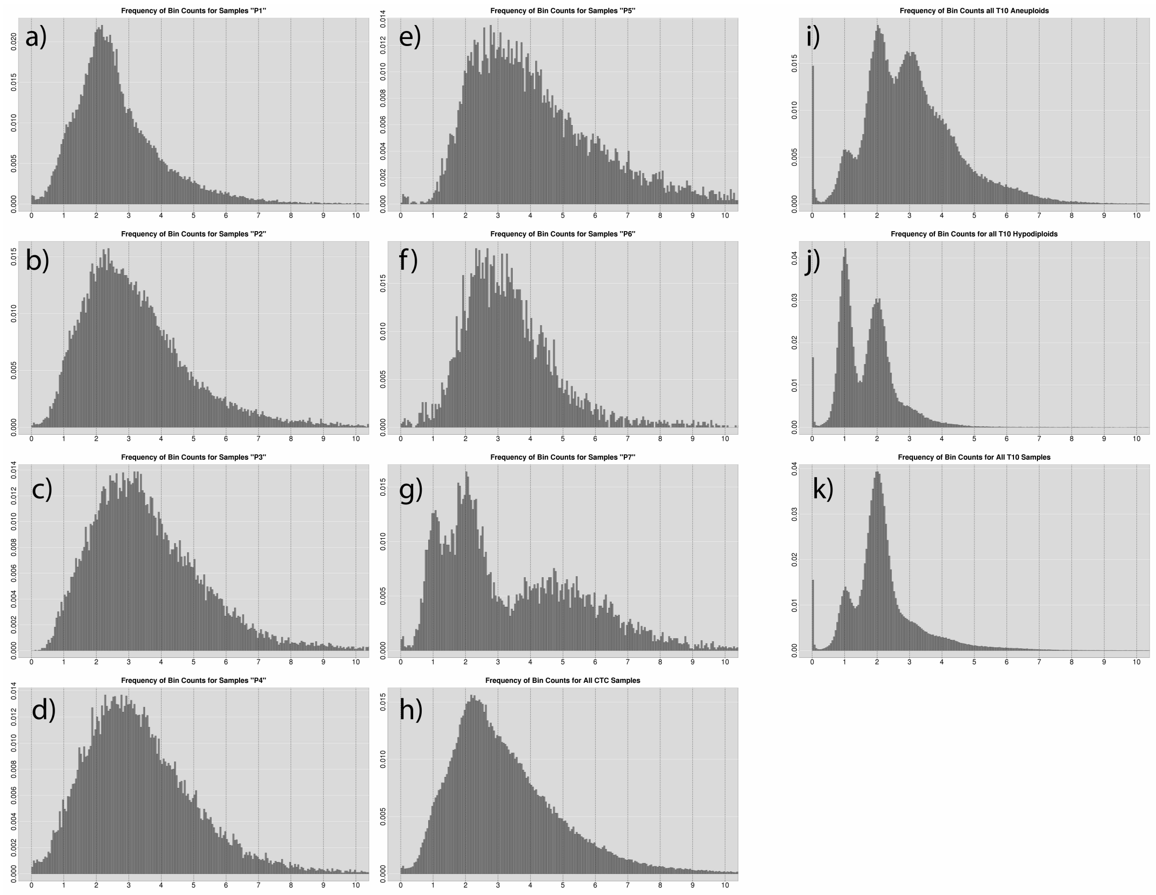

### Supplementary Figure 8: Histograms of the scaled bin counts

This analysis was performed across **(a-g)** patients 1-7 respectively, **(h)** all CTC patients, **(i)** all T10 hypodiploid samples, **(j)** all T10 aneuploid samples, and **(k)** all T10 samples.

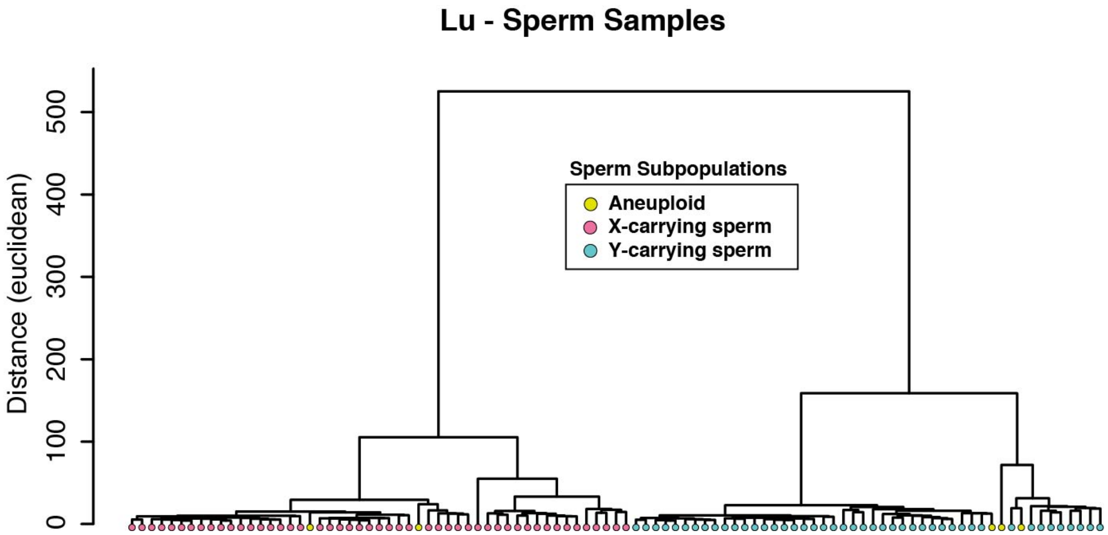

### Supplementary Figure 9: Ginkgo clusters the Lu *et al.* sperm samples

The major populations are defined by x- and y-carrying sperm. The original study identified 5 aneuploid cells (shown in yellow) with copy-number aberrations. Ginkgo is able to identify the same variants in the 5 aneuploid cells.
